# A bacterial secretosome for regulated envelope biogenesis and quality control

**DOI:** 10.1101/2022.01.12.476021

**Authors:** Daniel W. Watkins, Ian Collinson

## Abstract

As the first line of defence against antibiotics, the Gram-negative bacterial envelope and its biogenesis are of considerable interest to the microbiological and biomedical communities. All bacterial proteins are synthesised in the cytosol, so inner- and outer-membrane proteins, and periplasmic residents have to be transported to their final destinations *via* specialised protein machinery. The Sec translocon, a ubiquitous integral inner-membrane (IM) complex, is key to this process as the major gateway for protein transit from the cytosol to the cell envelope; this can be achieved during their translation, or afterwards. Proteins need to be directed to the inner-membrane (usually co-translational), otherwise SecA utilises ATP and the proton-motive-force (PMF) to drive proteins across the membrane post-translationally. These proteins are then picked up by chaperones for folding in the periplasm or delivered to the β-barrel assembly machinery (BAM) for incorporation into the outer-membrane. The core heterotrimeric SecYEG-complex forms the hub for an extensive network of interactions that regulate protein delivery and quality control. Here, we conduct a biochemical exploration of this ‘secretosome’ – a very large, versatile and inter-changeable assembly with the Sec-translocon at its core; featuring interactions that facilitate secretion (SecDF), inner- and outer-membrane protein insertion (respectively, YidC and BAM), protein folding and quality control (*e*.*g*. PpiD, YfgM and FtsH). We propose the dynamic interplay amongst these and other factors act to ensure efficient whole envelope biogenesis, regulated to accommodate the requirements of cell elongation and division. This organisation would be essential for cell wall biogenesis and remodelling and thus its perturbation would be a good strategy for the development of anti-microbials.

## INTRODUCTION

The bacterial envelope is essential for survival against an extraordinary range of physical and chemical environmental stresses. This protection has enabled their complete occupation of land and sea, including even the most inhospitable places and the invasion of more complex animal and plant hosts. While the envelope provides a barrier against the defence mechanisms deployed against bacteria, it is also an area of weakness for targeting by antibiotics and synthetic drugs.

The Gram-negative envelope is composed of a periplasm, containing a matrix of polymeric peptidoglycan (PG), sandwiched between inner- and outer-membranes. These membranes and intervening gelatinous space contain numerous proteins that need to be delivered, assembled and maintained at the right place (the inner-or outer-membrane, or periplasm), and time. Thus, the process is further complicated by the continuously changing state of the envelope, in response to changing environmental conditions and during cell division.

The classical view for this process involves the delivery of proteins to the inner-membrane – principally, the ubiquitous Sec machinery – for the delivery of proteins across or into it. Transport across the membrane, otherwise known as secretion, is usually achieved post-translationally by the cytosolic ATPase SecA in conjunction with the inner-membrane protein-channel complex SecYEG, facilitated by the ancillary sub-complex SecDFyajC [1] and driven by the proton motive force (PMF) [2–4]. By contrast, inner-membrane protein insertion occurs co-translationally, also *via* SecYEG, together with the highly conserved YidC [5, 6].

The SecYEG core-translocon, SecDFyajC and YidC combine to form a larger assembly known as the holo-translocon (HTL), capable of engaging in specialised post- and co-translational translocation [7, 8], and providing protein substrates with alternative routes across or into the inner-membrane. Proteins destined for the periplasm, outer-membrane or beyond that are recognised by the HTL and must travel through the centre of SecYEG, while α-helical membrane proteins slide through a lipid pool at the interface between SecYEG and YidC on their way to the bilayer of the inner-membrane [9].

Many of the proteins emerging into the periplasm are greeted by chaperones, such as SurA, Skp and DegP [10, 11], to facilitate folding, or degradation if they become mis-folded [12]. One class of such proteins are the outer-membrane proteins (OMPs) that primarily adopt a β-barrel transmembrane domain. These are delivered across the periplasmic space to the β-barrel assembly machinery (BAM) complex in the outer membrane for insertion and folding [13, 14]. Recently, we have shown that the HTL and BAM connect to form a trans-periplasmic inter-connection of the inner- and outer-membranes [15]. This most likely transient interaction provides a contiguous, unrestricted (*e*.*g*. by-passing the PG layer) pathway directly to BAM for outer membrane proteins (OMPs). Presumably this co-operation enables the required passage of very large quantities of OMPs, while protecting the transiting unfolded polypeptide from proteolysis and aggregation, for high maturation efficiency. The assembly seems to be stabilised by cardiolipin (CL) [15], a phospholipid required for the conferral of PMF stimulated secretion of OMPs [16].

Considering the knowledge presented above, we reasoned that there may be a hitherto unidentified trans-periplasmic complex, including the Sec and BAM complexes, plus several other permanent and guest residents, responsible for envelope biogenesis, quality control and cellular re-modelling. Such a complex may not have been distinguished before, *e*.*g*., by electron microscopy because the envelope is very crowed, or biochemically because of its instability and plasticity. The experiments described below explore the existence of such a *secretosome* and some (but certainly not all) of its prospective constituents and clientele, as well as their roles.

## RESULTS

### Very large membrane protein complexes of the *E. coli* envelope stabilised by cardiolipin

We began by analysing detergent extracts of total membranes of *E. coli*. Extracts were produced by solubilising membranes with the detergent dodecyl-maltoside (DDM) and subsequently fractionated by size exclusion chromatography. We know from previous findings that cardiolipin (CL) is required to stabilise the associated states of SecYEG-SecDFyajC-YidC (the holo-translocon) as well as the holo-translocon-BAM super-complex [17–19], so we reasoned that if an even larger Sec-associated complexes existed in the envelope then they too would be stabilised by CL. Thus, the size-exclusion experiments were conducted with and without augmented CL in the column buffer (0.02 % (w/v) DDM ± 0.002 % (w/v) CL). Interestingly, inclusion of CL during the chromatography did indeed enable the preservation of entities with very high apparent molecular weights of approximately 460 - 1700 kDa (calculated using calibration standards shown in Fig. S1, top panel), represented by a peak at 8.9 mL (Fig. 1a, light blue trace), which were diminished without the lipid supplement to form a peak at 10.2 mL (170 - 460 kDa) (Fig. 1a, dark blue trace). The average size of the mixed assemblies contained in both peaks were greater in size compared to purified SecYEG (Fig. 1a, magenta trace, peak at ∼160 kDa, true molecular weight = 74 kDa), HTL (Fig. 1a, red trace, peak at ∼200 kDa, true molecular weight = 249 kDa) and BAM (Fig. 1a, orange trace, peak at ∼300 kDa, true molecular weight = 209 kDa). Note the disparity between the apparent molecular weights of SecYEG, HTL and BAM and their true molecular weights, due to their annular detergent micelles and oligomerisation [20].

**Figure 1:**
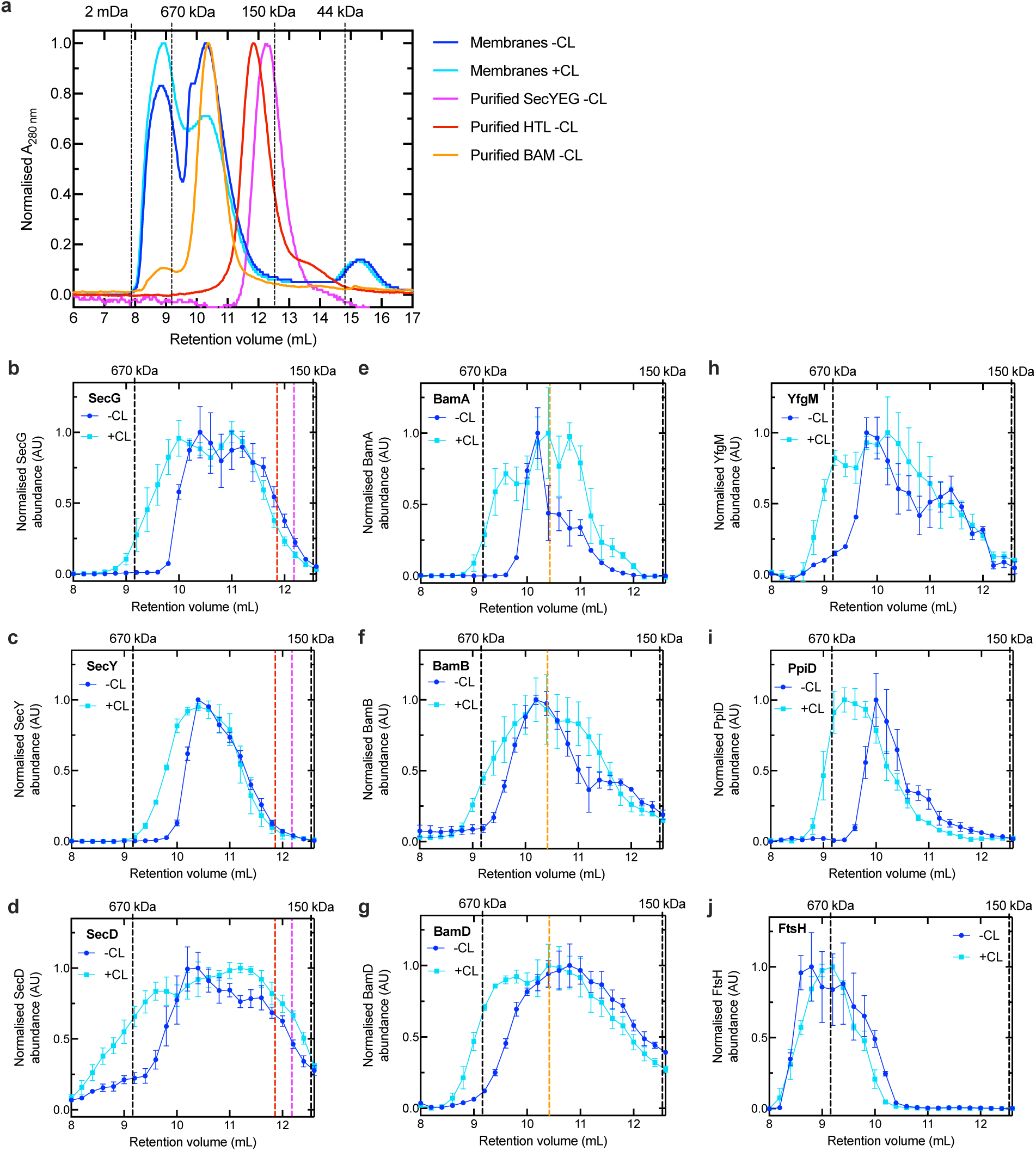
Size exclusion chromatography of *E. coli* total membranes. **(a)** Normalised A_280nm_ traces of solubilised *E. coli* total membranes purified on a S200 size exclusion column equilibrated in TSG with 0.02% DDM and with (+CL) or without (−CL) supplemented cardiolipin as indicated in the legend. Traces of purified SecYEG, BamA-E and HTL controls are also shown. Molecular mass standards were used to calibrate the column (Fig.S1) and their respective masses marked above the graph and indicated with dashed vertical lines. **(b-j)** Fractions from **(a)** were immunoblotted for proteins of interest (shown in the top left-hand corner of corresponding graphs). Blotting signal was quantified by densitometry and normalised to the maximum signal value. Both samples were analysed in triplicate (experimental repeats) and errors represent standard deviation. The peak of the elution profiles of the corresponding purified material from **(a)** are marked by magenta (SecYEG), red (HTL) and orange (BAM) dashed lines.

### A proportion of the envelope’s Sec translocons form part of a much larger assembly

The total *E. coli* membrane extract liberates a major early eluting peak, suggesting a high proportion of the cell’s total membrane proteins are contained within an assortment of large membrane protein complexes, with many of them stabilised by CL. Given the Sec translocon is formed of several proteins, and that it is known to interact with many others, we reasoned that the Sec subunits would be found in the large molecular weight peak. Therefore, we took the gel-filtrated fractions and looked for the individual proteins by immunoblotting. In the absence of CL, SecY and SecG of the core translocon and SecD of the HTL form similar elution profiles, mainly eluting between 10-12 mL (approximately 130 - 370 kDa, Fig. 1b-d, blue traces vs red and magenta dashed lines), corresponding to the low molecular weight peak seen in the chromatogram. In the presence of CL (Fig. 1b-d light blue trace), the elution profiles formed an additional shoulder between 9 - 10 mL (approximately 370-790 kDa), suggesting the formation of higher molecular mass complexes. In both the absence and presence of CL, the proteins elute at a higher apparent molecular mass than their purified counterparts (Fig. 1b-d, blue traces vs red and magenta dashed lines).

What other factors associate with SecYEG to account for this very large size difference? The most obvious are the HTL components SecDFyajC and YidC [7, 21], confirmed here by the coelution of SecD with SecY and SecG (Fig. 1b-d). However, the HTL alone (Fig. 1b-d, red trace) is not large enough to account for the early elution of the Sec subunits. Other factors must also be associated.

### Known Sec interactors form CL-dependent high molecular weight complexes

Next, we analysed the size-exclusion fractions for known interactors of the Sec-machinery. Further Western blots confirmed the presence of BAM (BamA, BamB and BamD) in the higher MW fractions, which was also dependent on CL (Fig. 1e-g), consistent with our previous findings [15]. The periplasmic chaperones YfgM and PpiD have both been shown to associate with one another and with the lateral gate of SecYEG for membrane protein departure [22, 23]. YfgM and PpiD were detected in the high MW peak along with the HTL and BAM (Fig. 1h, i). Once again, this associated state was preserved by CL. Finally, we looked for the inclusion of the AAA^+^ ATP-dependent protease FtsH – a cytosolic inner-membrane anchored quality control factor required for the degradation of misfolded proteins [24, 25] known to associate with SecY and YidC [26–29]. Interestingly, the protein was detected in the same higher MW fractions, but in this case independent of CL (Fig. 1j).

The overlapping, CL-dependent elution profiles of the Sec components and its known interactors points towards the presence of large complexes in the *E. coli* membrane. The existence of a ‘*secretosome*’ is a very intriguing prospect (discussed previously [15]). Thus, we set out to understand more about its stability and additional interactors.

### Removal of CL from the *secretosome* results in dissociation of its components

This secretosome is evidently prone to dissociation as its constituents were also detected in fractions corresponding to lower MWs (Fig. 1). This must be due, in part, to instabilities introduced by dissolution of the membranes and the extraction of specifically bound lipids – partially ameliorated by the augmentation of CL in the extraction buffer. When the high MW fractions were pooled and reapplied to the size exclusion column, the elution profiles of SecG, SecD and BamA volume were highly dependent on CL (Fig. 2). In the presence of CL the integrity of the higher MW complex containing the Sec and BAM complexes was maintained, while its omission resulted in a shift of all constituents towards the lower molecular weight regions of the chromatogram (higher volume). These experiments emphasise the importance of CL for the higher organisation of these and other factors.

**Figure 2:**
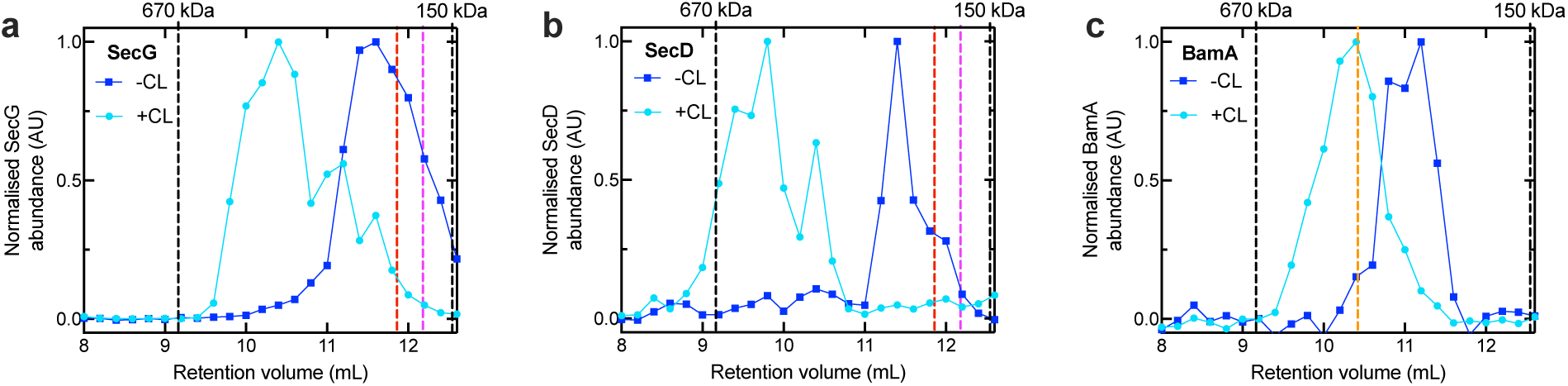
Stability of purified membrane complexes. Material collected between 9-10 mL after injection in Fig.1 were pooled and re-loaded onto the same column in TSG buffer with 0.02% DDM and with (+CL) or lacking (−CL) cardiolipin to analyse secretosome stability.

### Additional ancillary factors of the secretosome

In order to identify additional interaction partners, we utilised immune-purification and tandem mass tagging mass spectrometry (TMT-MS). In these experiments we immune-precipitated crude whole membrane extracts (± CL) with a monoclonal antibody raised against SecG – a non-essential constituent of the core-translocon SecYEG, and compared native *E. coli* membranes with those from a strain lacking SecG. Briefly, antibodies were incubated with the extracts which were then mixed with immobilised Protein-G resin. The resin was then stringently washed and the samples were prepared for TMT-MS. A *ΔsecG* strain enabled us to control for non-specific effects of CL (Fig.3a) and for non-specific binding to the antibody and resin (Fig. 3b-d). The three most significant interactors that were retained even without CL were: YajC – a component of the SecDFyajC sub-complex; YfgM – known to interact with SecYEG [22, 23] (Fig. 1h) and YiaD – a suppressor of defective temperature sensitive *bamD* with an affinity for peptidoglycan (Fig.3b). To the best of our knowledge, YiaD has not previously been identified as a Sec interactor.

**Figure 3:**
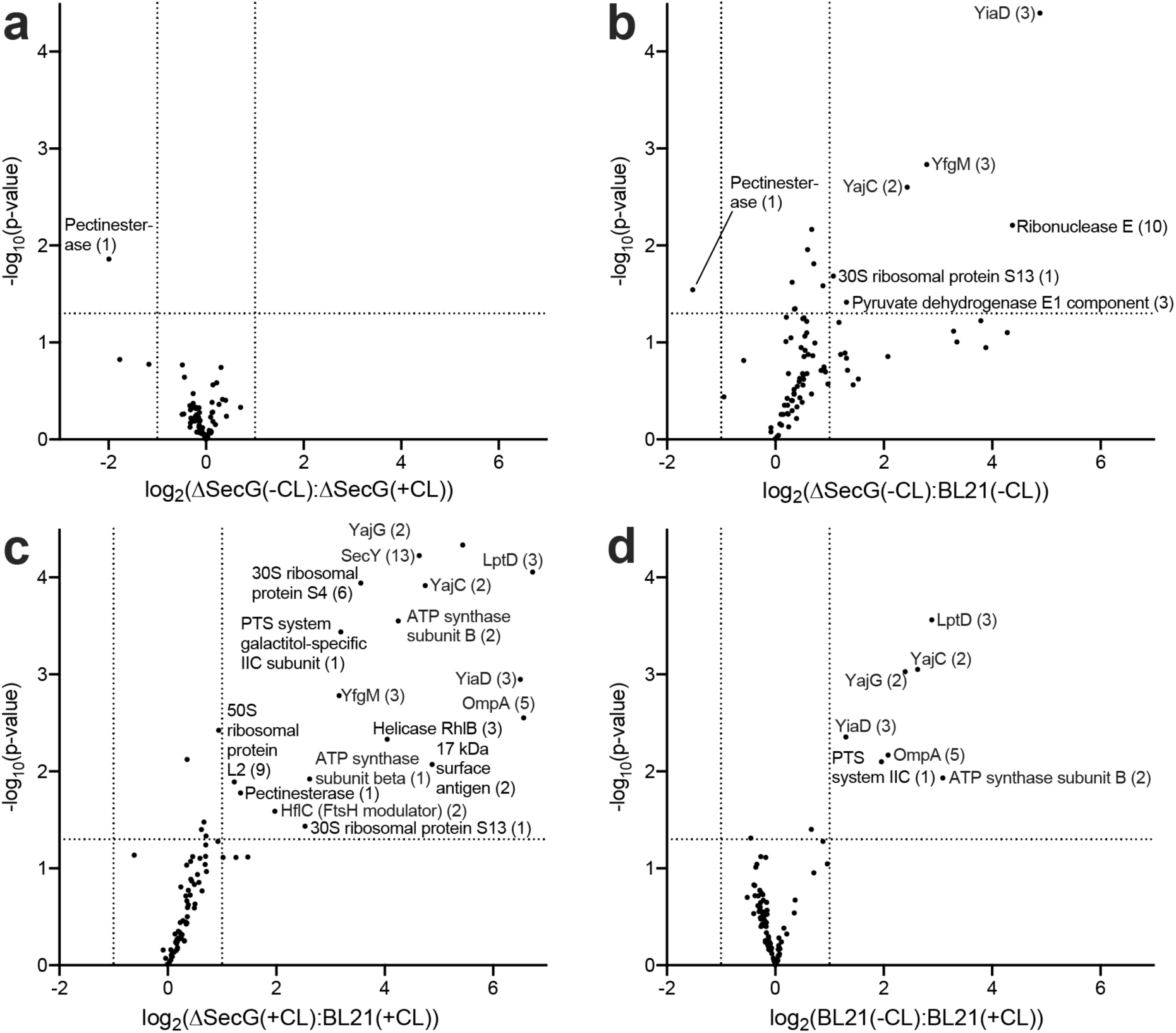
Tandem mass tag quantitative proteomic analysis of SecG co-IPs. Samples were prepared from SecG co-IPs of *E. coli* BL21 (DE3) and *E. coli* ΔSecG solubilised membranes in the presence or absence of supplemented cardiolipin, as indicated above respective graphs. X axis units represent fold change of protein abundance between the two described experiments. Each sample was prepared in triplicate (experimental repeats). An arbitrary cut off is applied at log2(fold change) = 1 and p-value = 0.05, both indicated by dotted lines for clarity. Proteins with significant abundance changes (p-value < 0.05 and log2(fold change) > 1) are annotated on graphs.

The changes resulting from the inclusion of CL were analysed by comparing results obtained from the ΔSecG and BL21 strains, both in the presence of CL (Fig.3c), and of the BL21 strain in the presence and absence of CL (Fig.3d). As expected, supplementing with CL resulted in more hits. YajC and YiaD were once again identified, suggesting a CL dependent interaction with the secretosome. Inclusion of CL also resulted in identification of SecY, likely due to the core translocon’s known dependence on CL for activity and stability. Interestingly, HflC was detected, which is a modulator of FtsH (see also Fig. 1j) and known interactor of YidC [29]. Two Sec substrates, OmpA and LptD, which are abundant β-barrel OMPs, were also detected. Their co-immuno-precipitation with SecG is likely due to capture during transit through SecY. The noted CL enhanced recovery of both OmpA and LptD clients, along with the putative secretosome constituents YajC and YiaD (Fig.3d), is suggestive of a CL-activated and stabilised complex. YajG was also identified when CL was retained in the extracts (Fig.3c,d), which intriguingly is an uncharacterised lipoprotein, and product of the *yajG-ampG* operon, the latter of which is important for the regulation of PG recycling and remodelling [30–32].

There were a few other identified proteins that require explanation. The ribosome subunits are abundant in cells and are known to interact with SecYEG. Pectinesterase was the only hit in the negative control experiment and was ignored as a result. The appearance of subunits of the ATP synthase is very curious, as a complex with SecYEG was previously identified in mixtures of membrane protein assemblies ejected from native *E. coli* membranes [33].

Inexplicable hits in proteomic-MS are commonplace. Some may have been identified due to being trapped in the secretosome. False negatives are also a feature as some membrane proteins resist proteolysis and/ or flight for MS. Moreover, extra caution was taken to ensure the protein G resin was stringently washed, and as a result proteins such as the BAM complex were detected, but not abundant enough to be accurately quantified.

### YfgM, PpiD and BAM interact with the HTL

The TMT-MS did not identify an anticipated enrichment of SecDF, BAM or PpiD. So, to verify their membership of the secretosome the SecG co-immunoprecipitants from the extracted native membranes were subjected to Western blotting (Fig. 4a,b). SecD was detected and further enriched by the inclusion of CL, most likely due to HTL stabilisation (Fig. 4a, middle). The same CL enrichment was observed for BamA, supporting previous results that the BAM complex interacts with HTL and not the SecYEG alone (Fig. 4, right) [7, 15]. The specificity of these interactions can be demonstrated through the analysis of membranes prepared from the *ΔsecG* strain, from which SecD and BamA were not co-immunoprecipitated (Fig. 4a, top). Similar experiments also recovered YfgM (highlighted also by TMT-MS; Fig. 3, c) and PpiD (Fig. 4b). In this case, the recovery was marginally increased in the presence of CL (Fig. 4c).

**Figure 4:**
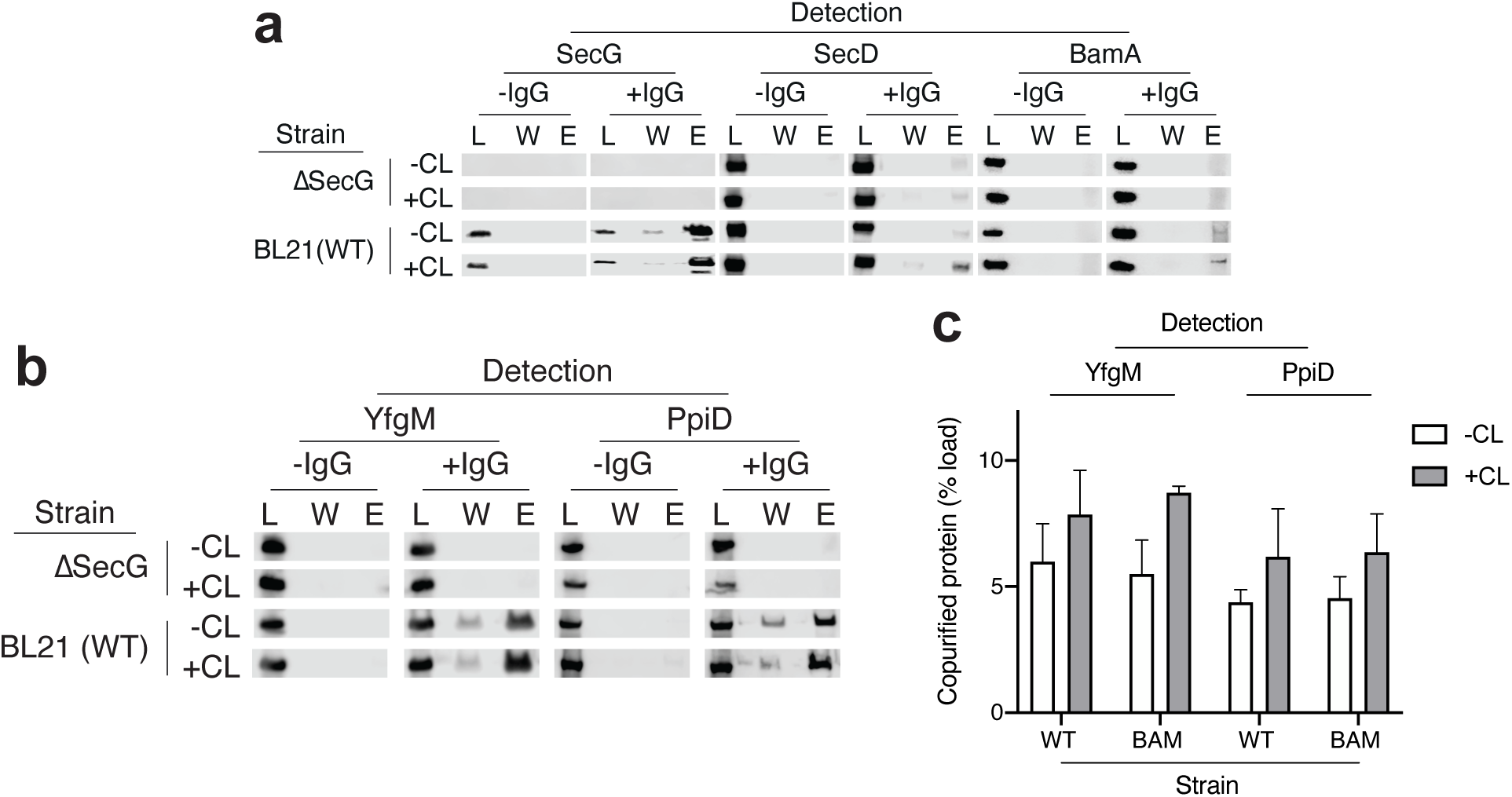
SecG co-immunoprecipitations. Co-IPs were conducted with membranes of *E. coli* BL21(DE3) or *E. coli* ΔSecG solubilised in 0.5% DDM and washed in the absence (−CL) or presence (+CL) of cardiolipin and in the absence (−IgG) or presence (+IgG) of SecG antibody. In (**a**) and (**b**) L = load (1% of total material), W = final wash (17% of total material) and E = eluate (17% of total material). Immunoblots are shown for SecG, SecD, BamA (**a**) and YfgM and PpiD (**b**). For (**b**) and (**c**), samples were prepared in triplicate (experimental repeats) and errors represent standard deviation.

## DISCUSSION

The experiments presented here all point to the existence of a very large assembly spanning the entirety of the Gram-negative cell envelope. The basic structures and function of this super-complex have already been discussed [15]. Beyond that we identify the involvement of additional components and speculate about the engagement of other factors for the formation of a ‘*secretosome*’: a dynamic and versatile hub for envelope biogenesis, quality control and remodelling (Fig. 5).

**Figure 5:**
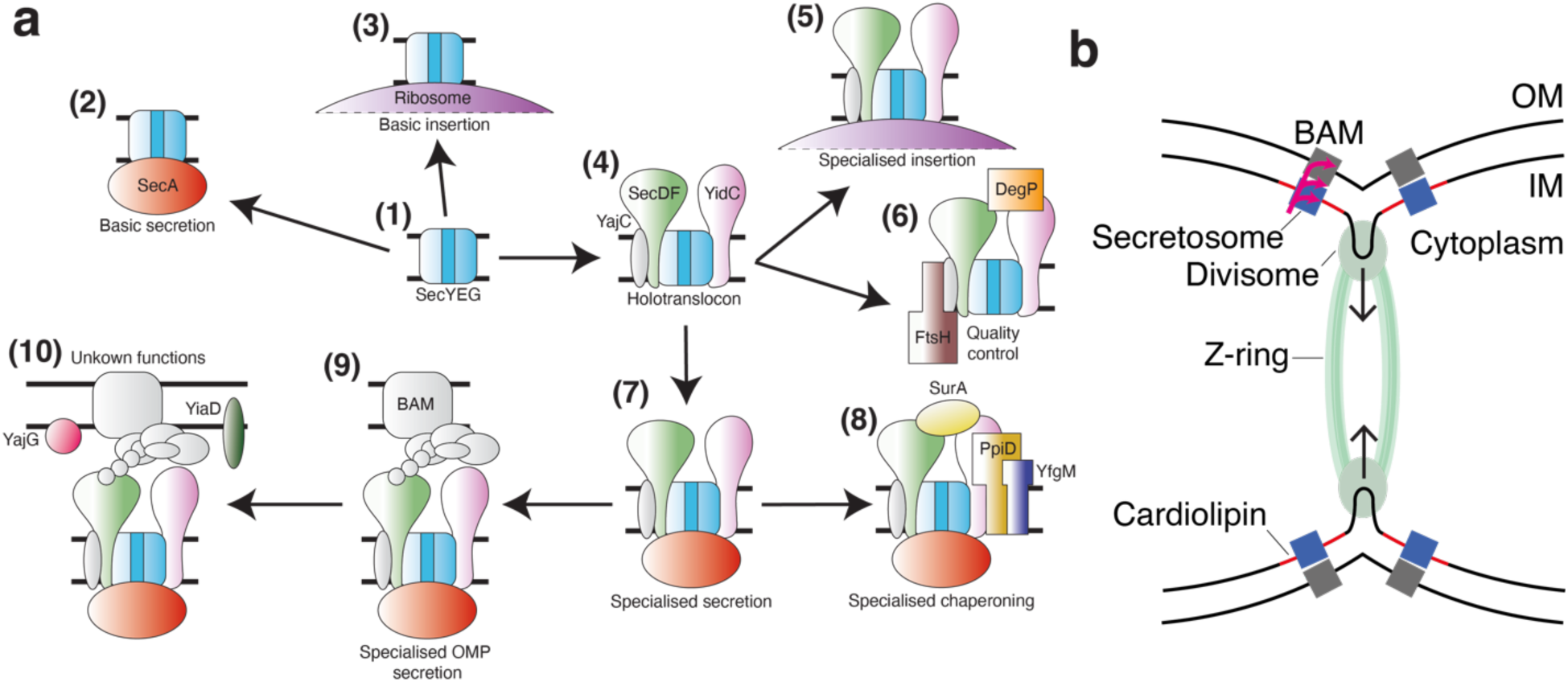
The Secretosome. **(a)** The core translocon, SecYEG (1), comes in many flavours, some of which are shown here. At the simplest possible level, secretion and inner membrane protein insertion can be conducted by SecYEGA (2) and ribsosome-SecYEG (3) respectively. On formation of the holotranslocon (HTL, 4), the secretosome functions, such as membrane protein insertion (5) and quality control enabled by FtsH (6), can become more specialised. As we have shown in previous work, HTL-SecA can conduct protein secretion (7) and is capable of binding inner membrane (PpiD and YfgM) and periplasmic chaperones (eg. SurA) (8), presumably to prevent aggregation of transiting proteins. HTL also interacts with BAM to facilitate specialised secretion of outer membrane proteins (9), and we have shown in this work that HTL interacts with the lipoproteins YiaD and YajG for unknown functions (10). Note that complexes shown here are not necessarily mutually exclusive. For example, chaperones or quality control proteins may bind to the insertion complexes or the HTL-BAM complex. **(b)** Recent studies have pointed towards localisation and direct interactions of components of the HTL and BAM with components of the divisome, where cardiolipin levels are elevated.

In this scenario, the basic activities of the core translocon SecYEG (Fig. 5a: 1) – protein secretion (Fig. 5a: 2) and membrane protein insertion (Fig. 5a: 3) – are streamlined, adapted or expanded by the association of various factors as required. The assembly of SecYEG with the ancillary components SecDFyajC and YidC forms the HTL (Fig. 5a: 4), which in all likelihood improves the efficiency of both core activities: YidC for membrane insertion (Fig. 5a: 5) and SecDF for secretion (Fig. 5a: 7) [5, 7, 34, 35]. Interestingly, a dynamic lipid pool at the interface between SecYEG and SecDFyajC-YidC is perfectly positioned to facilitate the former [9].

Chaperones also participate in the late stages of secretion, presumably in order to facilitate successful emergence and onward passage from the Sec-translocon, as well as folding and, where necessary, degradation. PpiD and YfgM, attached to the outer surface of the inner-membrane, and those of the periplasm, like Skp and SurA, presumably help facilitate the progression of proteins through and out of the secretosome (Fig. 5a: 8). While more work is needed to unravel the complex and dynamic interplay of the different chaperones at the secretosome, the different chaperones presumably have related, but subtly different activities. Indeed, SurA and Skp are at least partially redundant [10]. Perhaps the individuals, or various combinations of them, are required for different outcomes, such as delivery of β-barrelled OMPs to the BAM complex at the outer-membrane (Fig. 5a: 9), or early exiting of globular proteins into the periplasm.

Translocating proteins must on occasion become misfolded and/ or become trapped within the translocon [36]. This could irreversibly block the secretosome and potentially increase the conductance of small ions, compromising the energy conserving function of the inner membrane. In addition to the loss of functional export sites, this would prove catastrophic if left unresolved. Therefore, the conformational state of such a trapped complex might be recognised by factors, such as the chaperones mentioned above, whose recruitment could facilitate the clearance of denatured protein to nearby or associated proteases in the periplasm (*e*.*g*. DegP, FtsH; Fig. 5a: 6). Perhaps the secretosome can even be programmed for retro-translocation of such trapped substrates back through the holo-translocon for cytosolic degradation, for instance by the associated FtsH. Similarly, catastrophic blockages in the secretosome could be resolved by long-range conformational changes back to FtsH, for proteolytic degradation of SecY, and destruction of the defunct assembly. FtsH has indeed been shown previously to act in this way [28].

The action of CL on the stabilisation of the holo-translocon [7], the holo-translocon-BAM assembly [15] and the secretosome (shown here) is interesting because the lipid is known to localise in microdomains [37–40]. Thus, these CL-enriched islands could be sites for localisation of the translocon, by virtue of its affinity for this lipid over any other. These localised hubs could then recruit specific factors of the inner- and outer-membranes to form a bespoke secretosome. This idea resonates with a recently emerging concept of trans-envelope cross-talk for organisation and turnover of the residents of the inner- and outer-membranes [41, 42]. From the point of view of this analogy, it is striking that the outer-membrane islands involved in inter-membrane organisation are indeed enriched in BAM [42].

In this way we propose that specialised secretosomes localise, aided by inter-membrane communication, to form envelope biogenesis hot spots for the provision of the required proteins to the inner-membrane, periplasm and outer-membrane. The high concentrations of CL at these sites, important for secretosome assembly, would also serve to activate SecA – conferring high levels of ATP and PMF driven translocation through the secretosome [16, 17]. The localisation of CL at areas of high membrane curvature, including division sites and the poles [38, 40, 43], provides focal points for the localisation and activation of trans-envelope secretosome, establishing discrete sites for cell wall biogenesis. On this point, it has been shown that while most SecA is generally membrane associated and mobile, a significant proportion of SecA is immobile (∼25%) [44]. Perhaps the immobilised SecA is engaged with the secretosome at CL rich envelope biogenesis hubs. Indeed, we know that the affinity of SecA for the holo-translocon is higher where there is an abundance of CL [7]. Moreover, the formation of these static docking sites for SecA are dependent on the PMF [44], which ties in with the conferral of PMF stimulated protein translocation by CL [16], and the requirement of PMF for secretosome activity [15].

The observed broad elution profile of the secretosome constituents we see is presumably due to the existence of various assembled combinations of SecYEG, SecDF-YajC, YidC, BAM and others. This might reflect the situation in native membranes wherein there may be many different requirements at a given time and location. Different super-complexes, with different activities, may be recruited or assembled on site and tuned according to functional requirement, whether that be for delivery of proteins to the periplasm, inner-membrane, outer-membrane and/ or for regions in high demand of quality control. Presumably, the distribution of different complexes in the envelope will change during different stages of the cell cycle, which appears to be the case for the BAM complex [45]. Indeed, The HTL components YidC and SecG along with BAM were recently shown to localise at sites of cell division, adjacent to divisome proteins where they are presumably recruited to produce the high quantities of proteins required to enable cell division (Fig. 5b) [46]. This also aligns with a proteomics study demonstrating an interaction between SecY and the cell elongation/ division protein RodZ, suggesting the secretosome directly interacts with the divisome [47]. The same study picks up interactions of RodZ and FtsH with BamA. Elsewhere, work in another lab on another organism showed RodZ to interact with SecA and FtsH [48], supporting links between the division and secretion apparatus. Finally, the activity of BamA is supported by an interaction with DolP [49], which in the context of cell division is interesting because DolP is recruited to the mid-cell in the latter stages of this process [50].

The active recruitment of different factors to the core Sec-translocon is indeed an interesting concept for both eukaryotic and bacterial secretion systems. The Sec61-complex of the ER does so – forming a holo-translocon with additional components, for functions including membrane protein insertion and glycosylation [51, 52]. In this respect, the association of the Sec62/ Sec63 complex plays a role in the recruitment of Hsp70 homologue (BiP) of the ER-lumen to help drive efficient secretion and folding [53]. Interestingly, other more specialist functions have been attributed to the Sec62/ Sec63 pairing [54]. Ancillary components might interact transiently in order to support specific functional requirements, *e*.*g*. the ER membrane protein complex (EMC) to facilitate the insertion of more hydrophilic trans-membrane helices [55, 56].

Going back to the core features of the secretosome, *i*.*e*. for the delivery and maturation of proteins to the inner membrane, periplasm and outer-membrane, the trans-envelope association of the holo-translocon and BAM complexes brings together a number of interesting motifs, namely the tetratricopeptide repeat (TPR; YfgM, BamD, BepA) and WD40 β-propeller (BamB). Both motifs provide scaffolds for various other proteins to interact. The same can be said of the inter-membrane associated translocons of the mitochondrial outer-(TOM) and inner-membranes (TIM23). These TPR scaffolding sites could serve to recruit additional factors (*e*.*g*. in bacteria: SurA, Skp or DegP) necessary for transport, such as for retrieval into the periplasm for folding/ degradation, or to facilitate outer-membrane insertion. Perhaps even they could provide staging posts in the thoroughfare for transiting proteins.

## Materials and Methods

### Plasmids, strains and protein purification

*E. coli* strains BL21 (DE3), C43 or ΔsecG (KN425 (W3110 M25 ΔsecG::kan)) (a gift from Prof. Frank Duong) were used for all experiments. SecYEG and HTL were expressed and purified as described previously. Briefly, SecYEG was expressed in *E. coli* C43 from pBAD (Amp^r^), solubilised from the membrane fraction with 1.5% (w/v) n-dodecyl-β-D-maltopyranoside (DDM, GLYCON Biochemicals GmbSeH) in 20 mM Tris pH 7.5, 130 mM NaCl, 10% glycerol (TSG) and purified using nickel-immobilised metal affinity chromatography (IMAC) followed by size exclusion chromatography (SEC) with an appended anion exchange column [20]. The HTL was expressed using the pACEMBL system (Amp^r^, Kan^r^, Cm^r^) and purified the same as for SecYEG except with 0.002% (w/v) cardiolipin supplemented into IMAC and SEC buffers. The plasmid for BAM complex expression, pJH114-BamA-E (Amp^r^), was a gift from Prof. Harris Bernstein. Purification of BAM from *E. coli* C43 was performed as described previously [15].

### Production and solubilisation of membranes for co-immunoprecipitations and size exclusion chromatography

Precultures of *E. coli* BL21 (DE3) containing 100 mL 2xYT media were inoculated and grown overnight at 37°C and 200 RPM. The following morning, 10 mL of preculture was used to inoculate 1 L of 2xYT broth in a 2 L flask and the culture was incubated in the same conditions as described above. When an OD_600nm_ of 1.0 was achieved, cells were harvested by centrifugation (5000 xg, 10 minutes, 4°C) and resuspended in 20 mL of TSG. The sample was then passed through a cell disruptor (Constant Systems Ltd.) for lysis (2 passes at 25 kPSI). Membranes were clarified from the sample by centrifugation (160000 xg, 45 minutes, 4°C), resuspended to 120 mg/mL in TSG and stored at −80°C for future use. Solubilisation was performed by the addition of DDM to 0.5% (w/v). After 1 hour of gentle rocking at 4°C, insoluble material was removed by centrifugation (160000 xg, 45 minutes, 4°C) and the soluble fraction taken for further analysis.

### Size exclusion chromatography

A Superdex S200 10/300 GL size exclusion chromatography (SEC) column (Cytivia) was equilibrated in TSG buffer supplemented with 0.02% (w/v) DDM in the presence or absence of 0.002% (w/v) *E. coli* cardiolipin (Avanti Polar Lipids). With the flow rate set to 0.25 mL/min, proteins solubilised from 60 mg of wet membrane pellet as described above were injected through a 0.5 mL loop. 8 mL after injection, fractions of 250 uL were collected for approximately 10 mL. For SecYEG, HTL and BAM controls, approximately 350 µg of purified protein was loaded using the same conditions as for the solubulised membranes. Protein standards were obtained from Merck.

### Co-immunoprecipitations

For each reaction, 550 µL of solubilised membranes (66 mg of wet membrane pellet) were prepared as described above and 9 µL of purified monoclonal SecG antibody from our laboratory stocks was added. Samples were incubated overnight gently rocking at 4°C. 125 µL of Protein G Resin (Amintra) was prepared by washing 250 µL of the suspended resin in the manufacturer’s storage buffer in a microcentrifuge spin column 3 times with 500 µL of buffer containing 250 mM NaCl, 20 mM HEPES pH 8 (IP buffer) and centrifuge settings of 500 xg for 1 minute at 4°C. After the final wash, the resin was resuspended in 500 µL of IP buffer supplemented with 2% bovine serum albumin and incubated overnight in the same conditions as the membranes.

The following morning, the resin was washed using the same procedure as described above, but resuspended in 250 µL of IP buffer on the final step. 50 uL of material was removed from the solubilised membranes as a loading control and the resin and membranes were mixed and left to gently rock at 21°C for 3 hours. The resin was washed in a microcentrifuge spin column 6 times with 400 µL IP buffer containing 0.02% DDM in the presence or absence of 0.002% (w/v) *E. coli* cardiolipin. After the 6^th^ spin, the resin was resuspended in 150 µL of IP buffer with DDM ± cardiolipin and 50 µL of the suspended resin removed for proteomic analysis. The remaining liquid in the spin column was once again removed by centrifugation and this time collected in a fresh tube for analysis by SDS PAGE (last wash sample). Finally, bound proteins were removed from the resin by addition of 150 µL of 1x LDS sample buffer followed by centrifugation, again collecting the sample in a fresh tube.

### SDS PAGE and immunoblotting

SEC samples were analysed by SDS PAGE (NuPAGE™ 4 to 12%, Bis-Tris, 1.0 mm, Midi Protein Gel, 26-well) and transferred with a Power Blotter XL System (Invitrogen™) onto 0.45 µm nitrocellulose blotting paper (Cytivia Amersham™ Protran™). Immunoblotting was performed by incubating with either purified mouse antibody (SecG, SecY, both diluted 1/10000, from our laboratory stocks) or rabbit antiserum (SecD from our laboratory stocks, or BamA, BamC and BamD, a gift from Harris Bernstein, or a single antisera raised against both YfgM/PpiD, a gift from Prof. Daniel Daley, or FtsH, a gift from Prof. Joen Luirink, all diluted 1/5000), followed by incubation with anti-rabbit or anti-mouse HRP-conjugated secondary antibody (Life Technologies). A homemade ECL kit was used for imaging. Images were acquired for 10 minutes with an Odyssey-Fc imaging system (LI-COR Biosciences) and densitometry performed with the Image Studio Light software (LI-COR Biosciences). Graphs were produced with the Prism 8 software package.

### Tandem mass tag quantitative proteomic analysis of SecG co-IPs

Immuno-isolated samples were reduced (10 mM TCEP, 55°C for 1 h), alkylated (18.75 mM iodoacetamide, room temperature for 30 minutes) and then digested from the beads with trypsin (2.5 µg trypsin; 37°C, overnight). The resulting peptides were then labeled with TMT eleven-plex reagents according to the manufacturer’s protocol (Thermo Fisher Scientific, Loughborough, LE11 5RG, UK) and the labelled samples pooled and desalted using a SepPak cartridge according to the manufacturer’s instructions (Waters, Milford, Massachusetts, USA). Eluate from the SepPak cartridge was evaporated to dryness and resuspended in 1% formic acid prior to analysis by nano-LC MSMS using an Ultimate 3000 nano-LC system in line with an Orbitrap Fusion Tribrid mass spectrometer (Thermo Scientific).

In brief, peptides in 1% (v/v) formic acid were injected onto an Acclaim PepMap C18 nano-trap column (Thermo Scientific). After washing with 0.5% (v/v) acetonitrile 0.1% (vol/vol) formic acid peptides were resolved on a 250 mm × 75 μm Acclaim PepMap C18 reverse phase analytical column (Thermo Scientific) over a 150 min organic gradient, using 7 gradient segments (1-6% solvent B over 1 minute, 6-15% B over 58 minutes, 15-32% B over 58 minutes, 32-40% B over 5 minutes, 40-90% B over 1 minute, held at 90% B for 6 minutes and then reduced to 1% B over 1 minute) with a flow rate of 300 nL min^-1^. Solvent A was 0.1% formic acid and Solvent B was aqueous 80% acetonitrile in 0.1% formic acid. Peptides were ionized by nano-electrospray ionization at 2.0 kV using a stainless-steel emitter with an internal diameter of 30 μm (Thermo Scientific) and a capillary temperature of 275°C.

All spectra were acquired using an Orbitrap Fusion Tribrid mass spectrometer controlled by Xcalibur 3.0 software (Thermo Scientific) and operated in data-dependent acquisition mode using an SPS-MS3 workflow. FTMS1 spectra were collected at a resolution of 120 000, with an automatic gain control (AGC) target of 200 000 and a max injection time of 50 ms. Precursors were filtered with an intensity threshold of 5000, according to charge state (to include charge states 2-7) and with monoisotopic peak determination set to peptide. Previously interrogated precursors were excluded using a dynamic window (60 s +/-10 ppm). The MS2 precursors were isolated with a quadrupole isolation window of 1.2 m/z. ITMS2 spectra were collected with an AGC target of 10000, max injection time of 70 ms and CID collision energy of 35%.

For FTMS3 analysis, the Orbitrap was operated at 50 000 resolution with an AGC target of 50 000 and a max injection time of 105 ms. Precursors were fragmented by high energy collision dissociation (HCD) at a normalised collision energy of 60% to ensure maximal TMT reporter ion yield. Synchronous Precursor Selection (SPS) was enabled to include up to 5 MS2 fragment ions in the FTMS3 scan.

The raw data files were processed and quantified using Proteome Discoverer software v2.1 (Thermo Scientific) and searched against the UniProt *Escherichia coli* (strain B BL21-DE3) database (downloaded January 2020: 4172 entries) using the SEQUEST HT algorithm. Peptide precursor mass tolerance was set at 10 ppm, and MS/MS tolerance was set at 0.6 Da. Search criteria included oxidation of methionine (+15.995 Da), acetylation of the protein N-terminus (+42.011 Da) and Methionine loss plus acetylation of the protein N-terminus (−89.03 Da) as variable modifications and carbamidomethylation of cysteine (+57.021 Da) and the addition of the TMT mass tag (+229.163 Da) to peptide N-termini and lysine as fixed modifications. Searches were performed with full tryptic digestion and a maximum of 2 missed cleavages were allowed. The reverse database search option was enabled and all data was filtered to satisfy false discovery rate (FDR) of 5%.

## Acknowledgments

We are particularly grateful for the generosity of Dr Harris Bernstein for the kind gifts of the *bamABCDE* expression construct (pJH114) and antibodies. Thanks to Prof. Daniel Daley for the PpiD/YfgM antiserum and to Prof. Joen Luirink the FtsH antibody. Thanks to Kate Heesom of the University of Bristol Proteomics Facility for assistance with proteomics experiments.

## Author contribution

DWW and IC conceived and designed experiments; DWW conducted experiments; DWW and IC wrote the manuscript; IC secured funding and led the project.

## Declaration

the authors declare no competing interests.

## Data and materials availability

All data are available in the main text or the supplementary materials.

## SUPPLEMENTAL FIGURES

**Figure 1 supplement:**
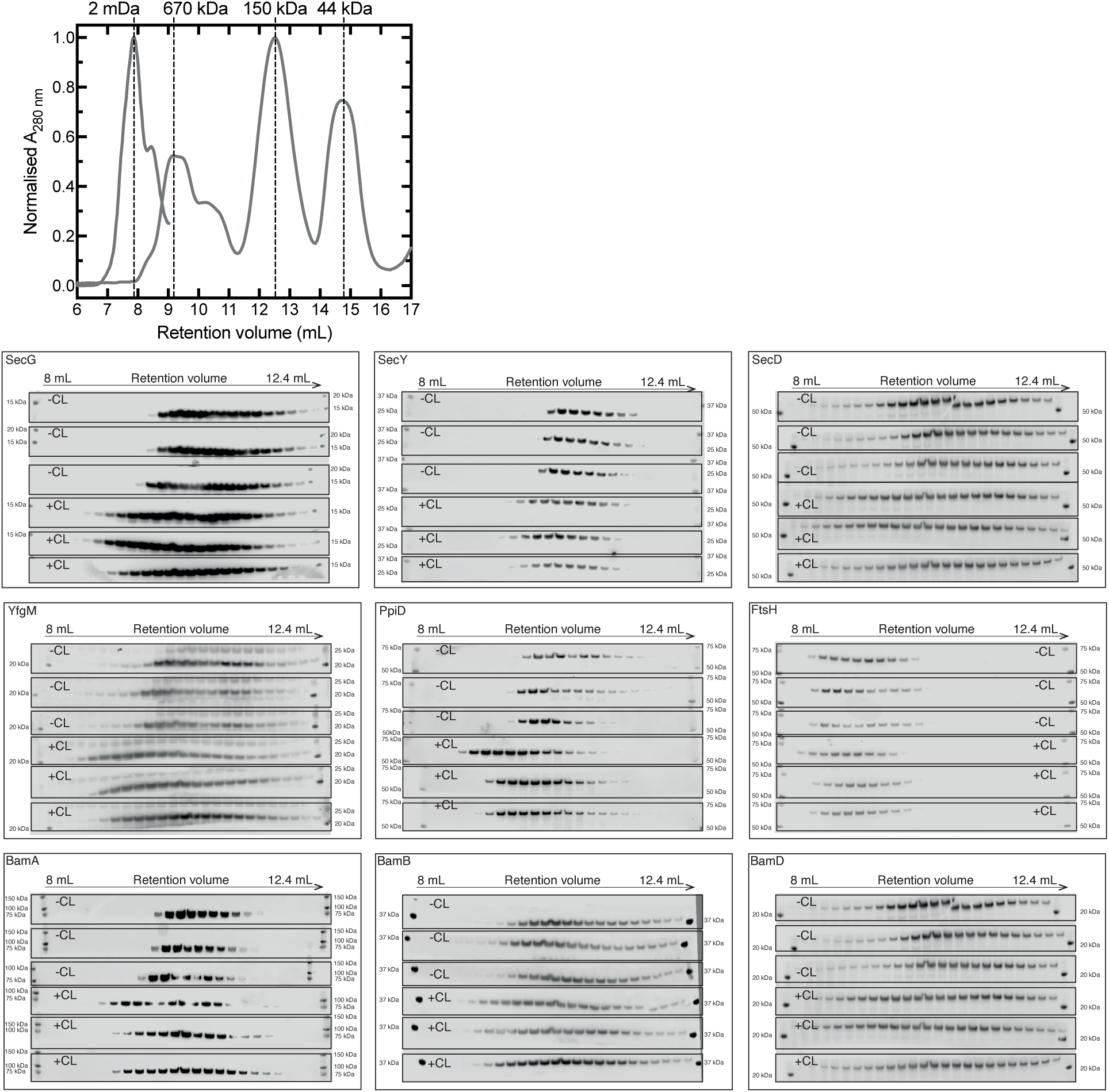
Immunoblots of S200 fractions of solubilised *E. coli* BL21 (DE3) membranes. Molecular mass protein standards (top) were used to calibrate the size exclusion column used in Fig.1. Densitometry was used to calculate relative signal abundance of immunoblots (bottom) and plotted against retention volume in Fig.1b-j.

**Figure 2 supplement:**
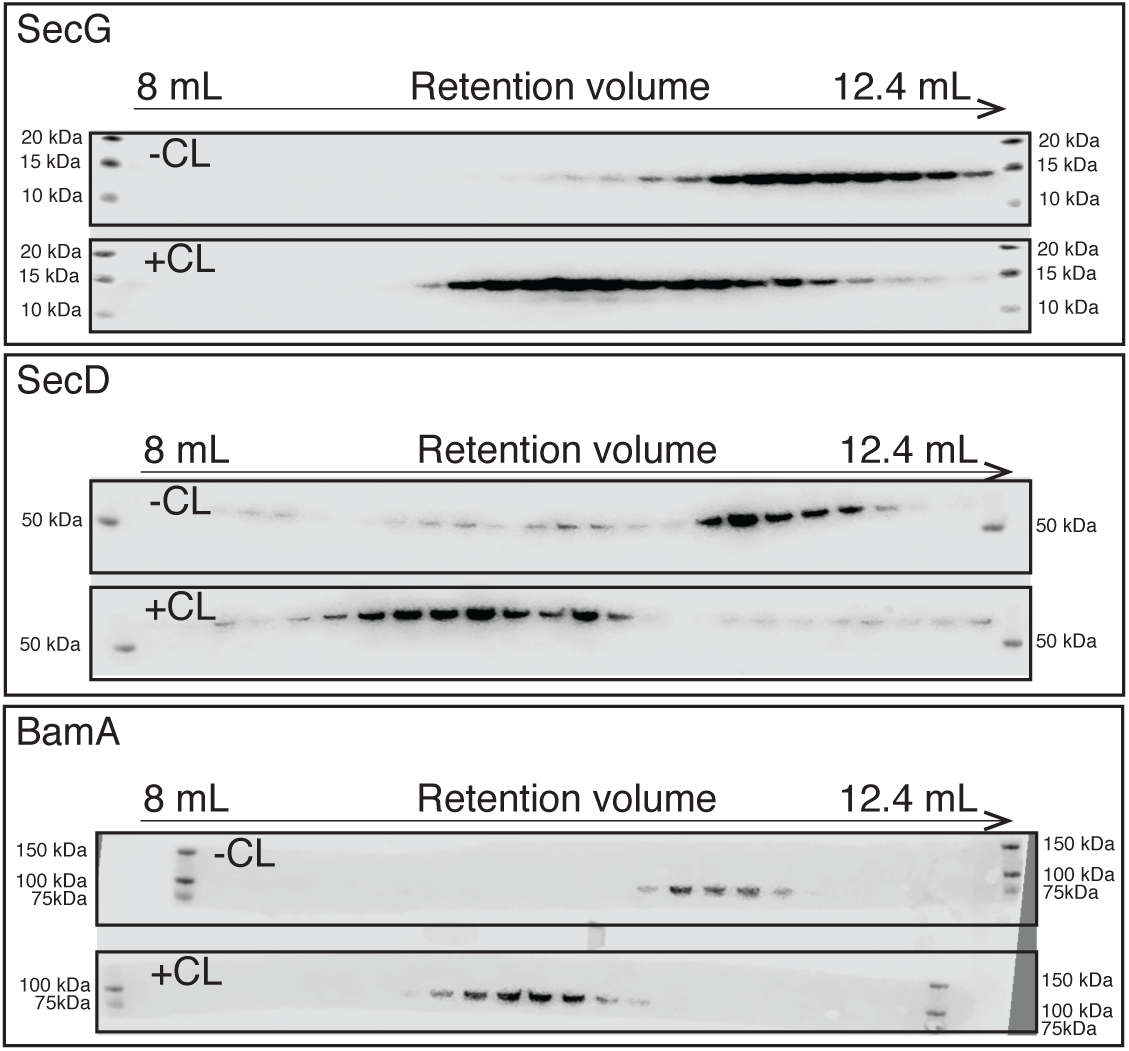
Immunoblots of S200 fractions of purified membrane protein complexes. Fractions collected between 9-10 mL after injection in Fig.1 were pooled and reloaded onto the same column in TSG buffer with 0.02% DDM and with (+CL) or lacking (−CL) cardiolipin to analyse secretosome stability. These raw blots were analysed by densitometry to give relative signal abundances shown in Fig.2.

**Figure 4 supplement:**
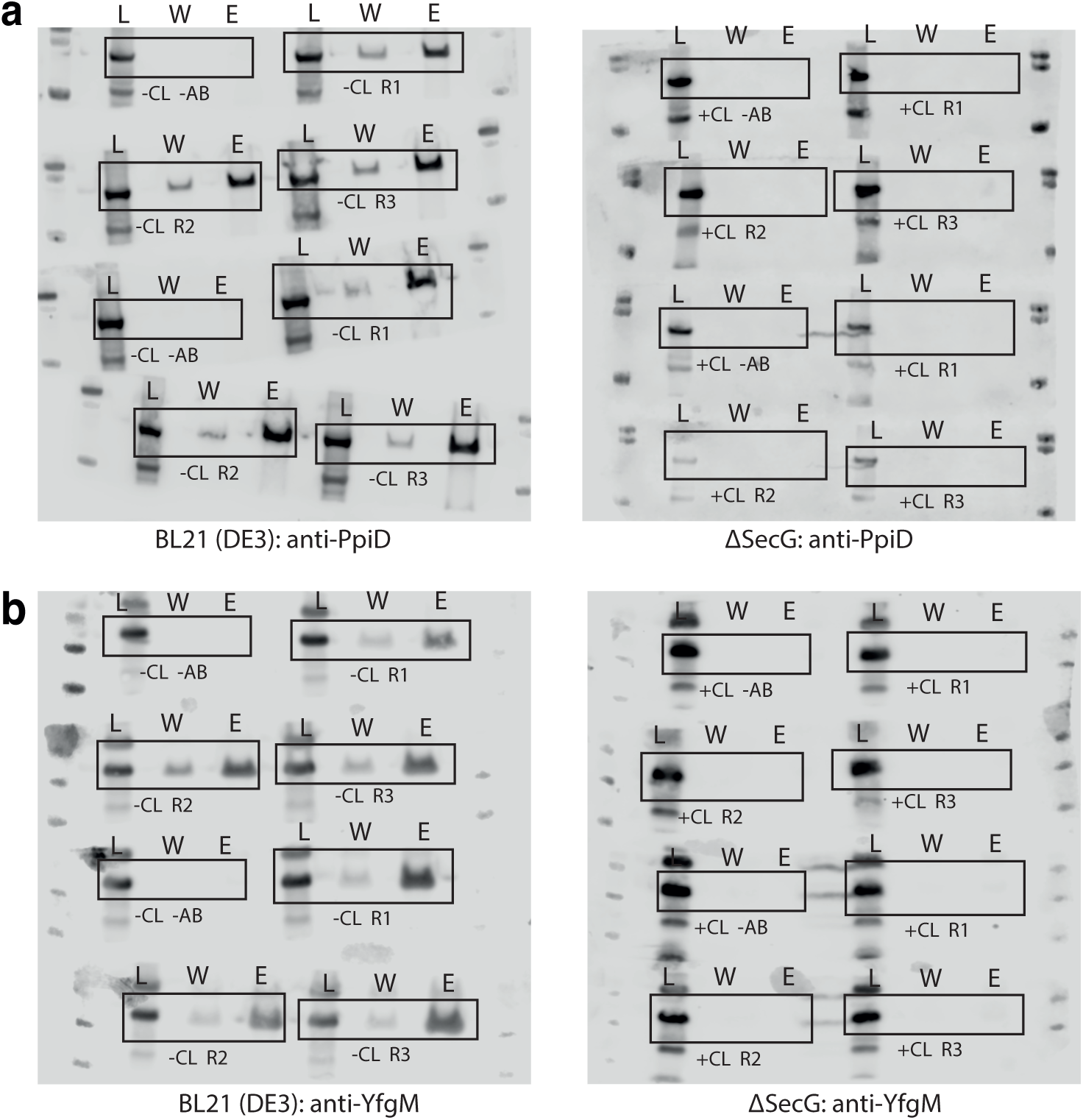
PpiD and YfgM raw western blots of SecG co-immunoprecipitations. Co-IPs were conducted with membranes of *E. coli* BL21(DE3) (left column) or *E. coli* ΔSecG (right column) solubilised in 0.5% DDM and washed in the absence (−CL) or presence (+CL) of cardiolipin and in the absence (-AB) or presence (+AB) of SecG antibody. L = load (1% of total material), W = final wash (17% of total material) and E = eluate (17% of total material). Samples were prepared in triplicate (experimental repeats, indicated by boxes marked ‘R1’, ‘R2’ and ‘R3’).

